# Evolutionary divergence of locomotion in two related vertebrate species

**DOI:** 10.1101/2021.02.11.430752

**Authors:** Gokul Rajan, Julie Lafaye, Martin Carbo-Tano, Karine Duroure, Giulia Faini, Dimitrii Tanese, Thomas Panier, Raphaël Candelier, Jörg Henninger, Ralf Britz, Benjamin Judkewitz, Christoph Gebhardt, Valentina Emiliani, Georges Debregeas, Claire Wyart, Filippo Del Bene

## Abstract

Locomotion exists in diverse forms in nature and is adapted to the environmental constraints of each species^1^. However, little is known about how closely related species with similar neuronal circuitry can evolve different navigational strategies to explore their environments. We established a powerful approach in comparative neuroethology to investigate evolution of neuronal circuits in vertebrates by comparing divergent swimming pattern of two closely related larval fish species, *Danionella translucida* (DT) and *Danio rerio* or zebrafish (ZF)^2,3^. During swimming, we demonstrate that DT utilizes lower half tail-beat frequency and amplitude to generate a slower and continuous swimming pattern when compared to the burst-and-glide swimming pattern in ZF. We found a high degree of conservation in the brain anatomy between the two species. However, we revealed that the activity of a higher motor region, referred here as the Mesencephalic Locomotion Maintenance Neurons (MLMN) correlates with the duration of swim events and differs strikingly between DT and ZF. Using holographic stimulation, we show that the activation of the MLMN is sufficient to increase the frequency and duration of swim events in ZF. Moreover, we propose two characteristics, availability of dissolved oxygen and timing of swim bladder inflation, which drive the observed differences in the swim pattern. Our findings uncover the neuronal circuit substrate underlying the evolutionary divergence of navigational strategies and how they are adapted to their respective environmental constraints.

## Main Text

*Danionella translucida* (DT) are minute cyprinid fish that show an extreme case of organism-wide progenesis or developmental truncation which leads to a small adult body size combined with a partially developed cranium without a skull roof. This feature together with its transparency throughout the adult stages makes them interesting for functional neuroscience studies allowing the imaging of the entire brain at cellular resolution^3,4^. However, studies on ossification in *Danionella sp.* demonstrate that most bones affected by truncation are formed later in the development of ZF^5,6^. Hence, in the early stages of their development, DT and ZF are highly comparable. DT and ZF are also found in similar freshwater environments in Asia and are evolutionarily very closely related^2,7^. This proximity is an advantage for comparative studies of larval DT and ZF as it provides a unique opportunity to understand how differences in behaviors can arise from relatively conserved neuronal circuits.

During undulatory swimming, animal experiences viscous and inertial forces in the fluid. Based on the body length, the hydrodynamics dictating the swimming also changes^8,9^. However, the body length of ZF and DT falls in a similar range of few millimeters which leads to a transitional flow regime for both (Fig. 1a-b; size range: 4.1 to 4.9 mm)^8^. To compare the kinematics of their spontaneous swimming, we recorded their movement with a high-speed imaging system. Larval ZF are known to swim in a beat-and-glide pattern wherein a burst of tail activity lasting ~140 ms enables them to swim at high speed and is followed by a passive glide phase^10^. In contrast, larval DT move at low speed by continuously beating their tail for few tens or even hundreds of seconds (Fig. 1c-d, Extended Data Movies 1 and 2). To compare the fine swimming kinematics of DT and ZF, we defined kinematic parameters based on half-tail beats, a unit common to the swimming pattern of the two species. (Extended Data Fig. 1). The continuous slow swims of larval DT occur with a smaller half tail beat frequency and a smaller maximum tail angle compared to larval ZF (Fig. 1e). In head-embedded preparation, continuous and intermittent swimming patterns were observed as well in larval DT and ZF, respectively (Fig. 1f): DT swims for 98.5 % of the total recording time compared to only 2% in ZF (Fig. 1g).

**Fig. 1:**
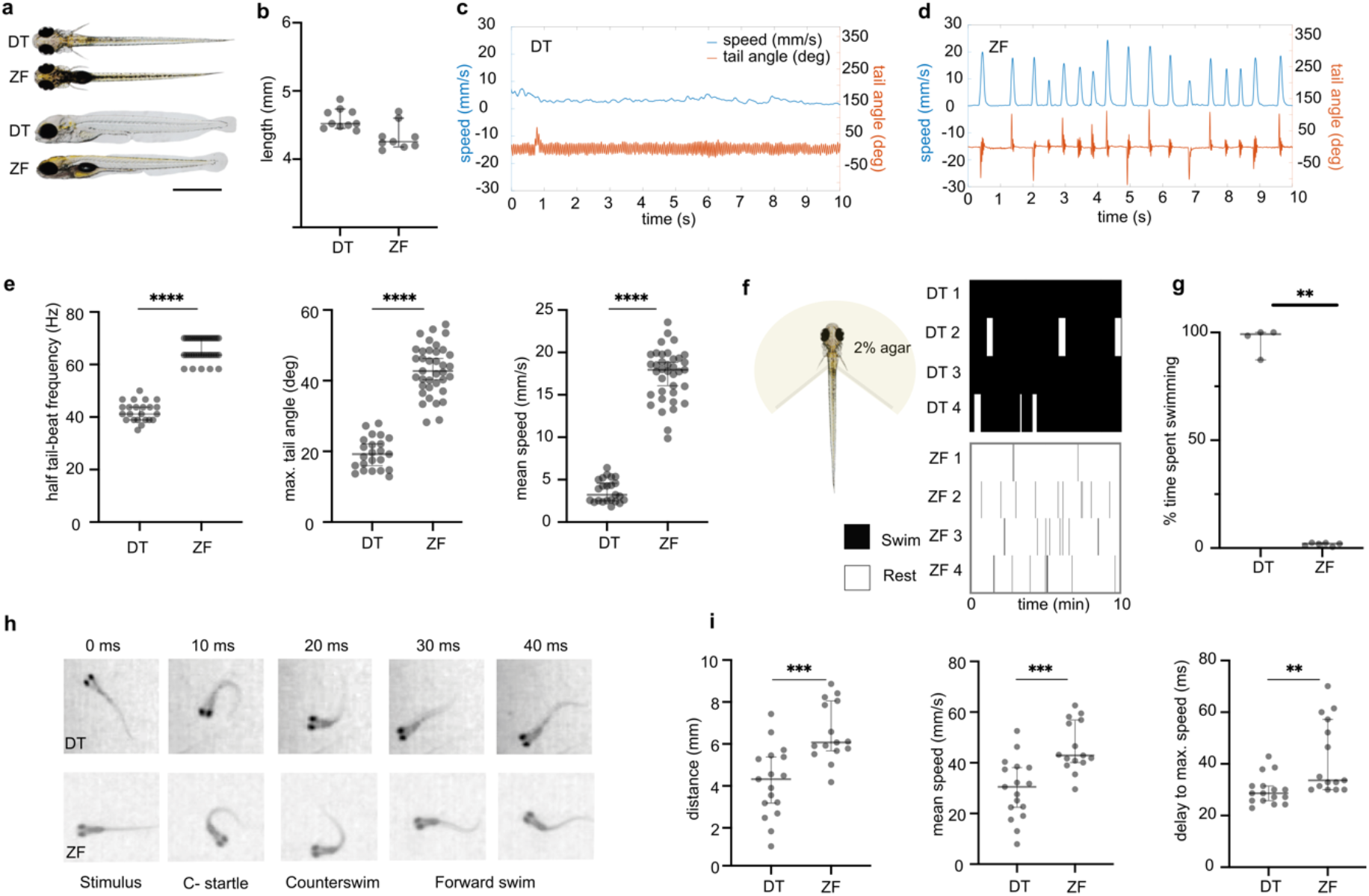
Kinematics of spontaneous swimming, head-embedded swimming and escape response in DT and ZF. Larval DT and ZF measure similar in size at 5 dpf. (a) A dorsal and lateral view of DT and ZF at 5 dpf is shown. (b) Measurement of body length in 5 dpf DT and ZF falls in a range of 4.1 to 4.9 mm. (N= 10 DT; N= 9 ZF). (c-d) A comparison of swimming pattern in 6 dpf DT and ZF. It demonstrates the continuous swimming pattern in DT with lower speed (cyan) and smaller tail angle (orange) when compared to the faster discrete swimming in ZF. (e) Swimming kinematics of DT and ZF in a spontaneous swimming assay. DT utilizes lower half tail beat frequency (Hz) and lower maximum tail angles (degree) to achieve swimming at lower speeds (mm/s) when compared to ZF (N= 23 DT, n = 494628 half tail beats and N= 37 ZF, n = 202176 half tail beats). (f) Tail movements in head-embedded preparations depicted in raster plots illustrate the prolonged swims of DT (top) compared to the short bouts of ZF (bottom). (g) The fraction of time spent actively swimming (% of total acquisition time) is higher in DT compared to ZF (N=5 DT and N= 6 ZF). (h) Qualitatively, the escape response after a tap stimulus is highly similar between DT and ZF. The images were acquired at 100 Hz. (i) Although DT covers a shorter distance at a lower mean speed, the time to achieve the maximum speed is lower in DT compared to ZF (DT: N=19 fish, n=141 events; ZF: N=15 fish, n=159 events). ** p<0.01, *** p=0.001, **** p<0.0001, Mann-Whitney test. All error bars show 95 % confidence interval.

In order to test the ability of larval DT to achieve fast speeds following sensory stimulation, we examined their escape response using tap-induced escape assay (Extended Data Movie 3). Fig. 1h shows the striking similarity in the escape response between DT and ZF. Both fish species initiate a fast C-bend followed by a counter bend. DT was found to swim with a lower mean speed and cover a smaller distance during this period compared to ZF (Fig. 1i, Extended Data Fig. 2). On the other hand, the delay to achieve maximum speed during escape was surprisingly smaller in DT. This faster response may compensate for the relatively lower speed of DT during an escape response (Fig. 1i). Our data shows that DT are capable of executing fast swims to escape, but have evolved a slow and continuous swimming mode during spontaneous exploratory behavior.

We further investigated how the distinct modes of spontaneous navigation shown by both species may impact their long-term exploratory kinematics. We monitored the swim trajectories of DT and ZF in a 35 mm diameter Petri dishes as shown in Fig. 2a-b. We then computed the mean square displacement (MSD) that quantifies the area explored by the animal over a given period of time (Fig. 2c). Surprisingly, although the DT mean forward velocity is significantly lower than ZF, the MSDs are comparable. This can be understood by considering differences in heading persistence in both species. At short-time scale, the larvae tend to swim in straight lines such that their trajectories can be considered ballistic. On longer time scales, reorientation events cumulatively randomize the heading direction and the dynamics becomes diffusive-like. The ballistic-to-diffusive transition time can be estimated by computing the decorrelation in heading direction, as shown in Fig. 2d. These graphs reveal a faster randomization of heading direction in ZF compared to DT. In ZF, the decorrelation function R(t) drops down to 0.3 in ~1 second, i.e. the typical inter-bout interval, then decays to 0 in 6-7 seconds. In DT, R(t) shows a small initial drop (down to 0.8) then slowly decays to 0 over the next ~8 seconds. The small initial decay in the first ~0.5 seconds can be interpreted as reflecting the short time-scale fluctuations in heading direction during run periods. The further slow decorrelation in turn results from the successive reorientation events that separate the periods of straight swimming, a process reminiscent of the classical run-and-tumble mechanism of motile bacteria^11,12^. The time-scale of this slow decay is expected to be controlled by the interval between successive reorientation events, which is of order of 5-10 seconds.

**Fig. 2:**
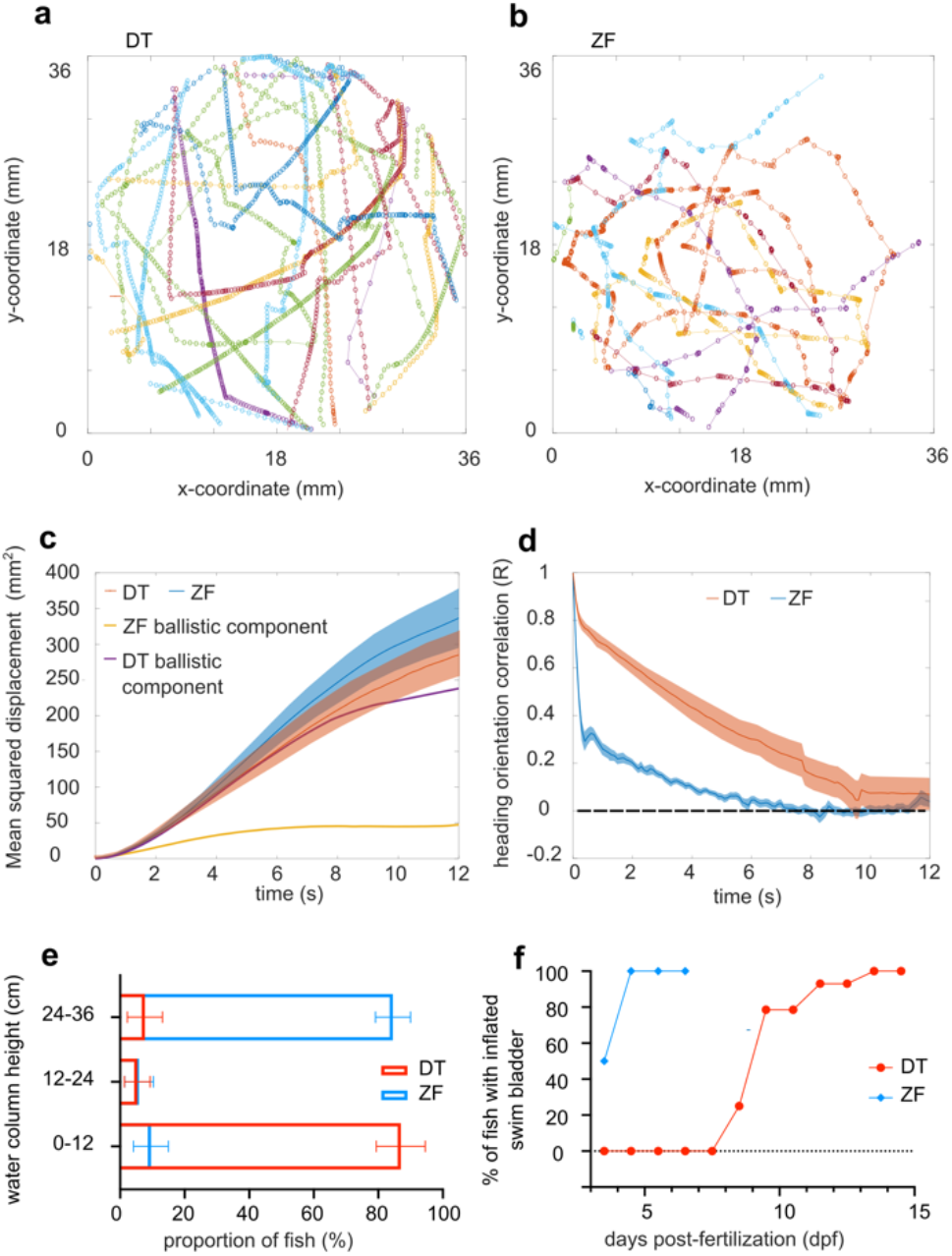
Long-term exploratory kinematics of DT and ZF. (a) and (b) represent a few tens of swim trajectories depicted in different colors from a single DT and ZF larva, respectively. (c) Mean squared displacement (MSD) in DT and ZF over time. The MSD over time for the ballistic component of DT and ZF is also overlapped on the plot. The ballistic component of the MSD was estimated as: 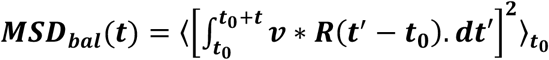. The error bars show s.e.m. (d) Decorrelation in heading persistence over time. R=1 indicates a perfect persistence in head direction whereas R=0 corresponds to a dull randomization. In ZF, R drops rapidly whereas this drop happens over longer period of time in DT. Hence, exploratory swimming in DT has a longer ballistic phase. The error bars show s.e.m. (e) DT larvae occupy the bottom of the water column whereas ZF larvae occupy the top. (N= ~30 fish and n = 10 readings in 3 replicates for each fish species). The error bars show 95% confidence interval. (f) Swim bladder inflation in ZF occurs earlier than in DT. The inflation of swim bladder in ZF occurs by ~ 4-5 dpf whereas DT inflates their swim bladder between ~10 and ~15 dpf. Sampled from a growing population of approximately N = 30 DT and 30 ZF.

To quantitatively assess the relative contribution of the ballistic vs diffusive components of the MSD over time, we estimated the former as:

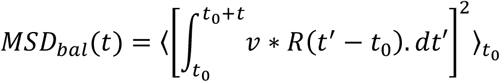

where *υ* is the mean instantaneous velocity. As expected, for a purely ballistic process, R=1 and *MSD_bal_ = (υt)^2^*, whereas for a purely diffusive process, R=0 and *MSD_bal_* = 0. For DT, this quantity correctly captures the MSD up to ~6 seconds (Fig. 2c), indicating that the ballistic component is dominant over this long initial period. In contrast, for ZF, the MSD departs from the ballistic component from 1 second onwards, i.e. after 1-2 bouts. In summary, our analysis shows that the pattern of navigation adopted by DT yields longer heading persistence, which almost exactly compensates for its intrinsically lower swimming speed and results in comparable long-term spatial explorations.

We next asked what selective environmental and physiological pressures might have led to the differing swimming patterns. At the environmental level, we explored the role of dissolved oxygen on these differences. During our field study in Myanmar, we found that adult DT were most abundant at the lower water levels of a small stream at ~50 cm, characterized by lower oxygen levels compared to the surface (O_2_= 8.75 mg/L at the surface; O_2_= 3.5 mg/L at 50 cm; O_2_= 2.8 mg/L at 80 cm) (Extended Data Fig. 3a). In contrast, adult ZF are reported in waters with variable DO concentration with a median of 5.55 ±1.64 mg/L but this lacks information on the depth at which it was recorded^13^. In the laboratory, we tested the occupancy of larval DT and ZF in a tall water column of 36 cm height. Larval DT were found to occupy the lower zone of the water column whereas larval ZF were found in the upper zone of the water column (Fig. 2e). This is consistent with previous observations that ZF adults spawn in very shallow environments while adult DT spawn in the narrow spaces in the bottom of the river bed^7,3^. As we have seen, upper layers of a water column in the wild are richer in dissolved oxygen (DO) when compared to the bottom layers^14,15^. Hence, the apparent preference of DT for deeper water would accompany a lower availability of DO in the wild. This lower DO availability would have consequences on swimming at two levels: during locomotion and at rest. During locomotion, a lower DO would act as a constraint to the maximum swimming speed that can be achieved by an animal (Extended Data Fig. 3b)^16^. At rest, it has been shown both, experimentally and analytically, that increased body movements with reduced stationary periods would be beneficial for larval fish in a low DO environment to be able to replenish DO in its immediate surrounding. (Extended Data Fig. 2c)^17,18^.

At the physiological level, we propose that a difference in the timing of swim bladder inflation in DT and ZF might have an important role in the differences that we observed in the swimming pattern. Consistent with previous reports, we observed inflation of swim bladder in ZF by ~4-5 dpf, whereas in DT population this occurred later between 10 and 15 dpf (Fig. 2f). As has been shown previously, without a swim bladder to regulate its buoyancy, a fish might need to continuously swim and exert a downward force to actively maintain its position in the water column^19^. This would also promote a continuous movement as observed in DT. However, it is important to note that the delayed inflation of swim bladder alone cannot explain the difference in swimming pattern. Indeed, we evaluated swimming in a population of DT and ZF larvae at 15 dpf, all with inflated swim bladders. Although a decrease was seen in the proportion of time spent swimming in 15 dpf DT when compared to 6 dpf DT, this value remained very high compared to ZF at 6 and 15 dpf (Extended Data Fig. 3d). Altogether, our data show that the slow and continuous swimming pattern of DT might be a result of a combination of factors including lower availability of DO and delayed inflation of swim bladder.

To determine the cellular underpinnings of the difference in swimming modes (continuous versus discrete) in the two species, we first investigated the organization of neuronal populations in DT. We initially examined reticulospinal neurons in the brainstem as these neurons projecting from hindbrain to spinal cord are known to play a role in locomotion control in all vertebrates^20,21,22^. We identified numerous cells of the mesencephalic nucleus of the medial longitudinal fascicle (nucMLF/nMLF), the rhombocephalic reticular formation (nucRE) and the rhombocephalic vestibular nucleus (nucVE) whose location of soma and morphology showed a high homology with cells previously described in ZF (Fig. 3b)^23,24^. For instance, we observed dendrites crossing the midline from the MeM cells (of nMLF) in DT as previously described in ZF (Fig. 3c). Next, we investigated the distribution of excitatory (glutamatergic) and inhibitory (glycinergic and GABAergic) neurons in the hindbrain. In ZF, these populations are described to be spatially organized in distinct stripes ordered according to their developmental age and neurotransmitter identity^25,26,27^. Consistent with this arrangement, the distribution of excitatory and inhibitory neurons in DT hindbrain also forms rostro-caudally running stripes (Fig. 3d-f, and cross-section in Extended Data Fig. 4). The nearly identical location of reticulospinal neurons, and the similar distribution of neuronal population points to a closely conserved bauplan of the hindbrain locomotor control region. However, homologous neurons could have different functions, as shown for neurons involved in feeding behaviors in nematodes^28^.

**Fig. 3:**
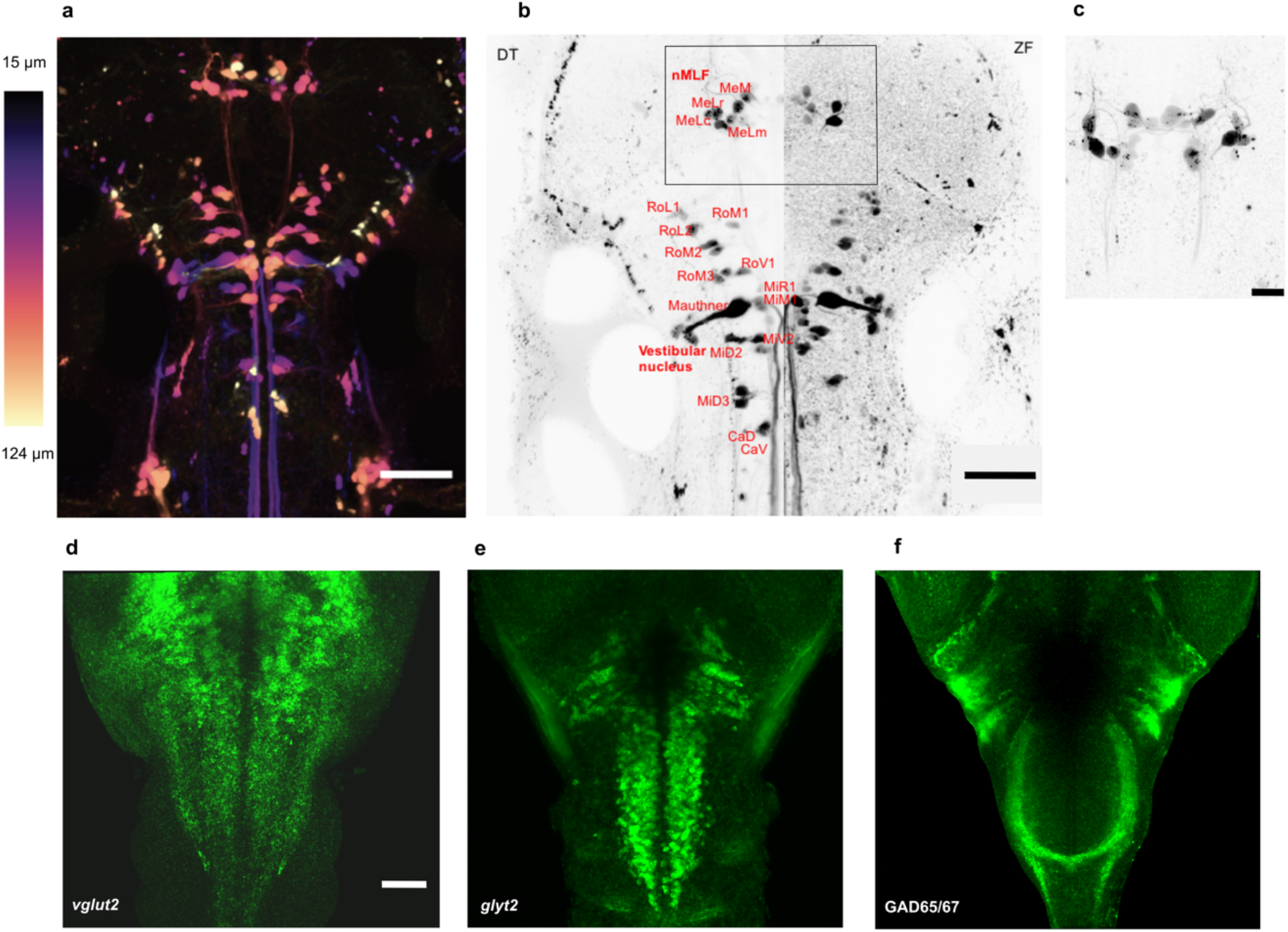
Distribution of reticulospinal neurons and excitatory and inhibitory neuronal cell types in the hindbrain of DT and ZF. (a) Distribution of reticulospinal (RS) neurons in the brainstem of DT. Maximum intensity is color coded for depth. Scale bar is 100 μm. (b) Comparison of RS neurons in DT and ZF. A cell-to-cell comparison of RS neurons in a maximum intensity projection of RS neurons in DT and ZF. The RS neurons in DT are annotated based on the description of RS neurons in ZF.^23,24^ A high degree of conservation is observed. Scale bar is 100 μm. (c) A closer look at the rectangular ROI from (b) in another DT fish. Maximum intensity projection of the nucleus of the medial longitudinal fasciculus (nMLF) in another DT fish is shown. Contralateral dendritic projections are observed in DT as noted in ZF^23^. Scale bar is 30 μm. Distribution of (d) glutamatergic (e) glycinergic and (f) GABAergic neurons in the hindbrain of DT. Performed using *in-situ* hybridization (ISH) and immunohistochemistry (IHC). Rostrocaudally running striped pattern of these neuronal types is observed in DT as noted in ZF before^25^. Anti-*vglut2a* + *anti-vglut2b* ISH and *anti-glyt2* ISH in (d) and (e), respectively. Anti-GAD65/67 IHC in (f). All images are a maximum intensity projection. Scale bar is 50 μm.

In order to identify functional differences in neuronal activation during spontaneous locomotion, we investigated the recruitment of neurons throughout the brain using whole-brain calcium imaging in larval DT and ZF. We generated a novel transgenic line *Tg(elavl3:H2B-GCaMP6s)* in DT where the calcium indicator, GCaMP6s was nuclear-targeted and expressed under a pan-neuronal promoter as previously done in ZF (Extended Data Fig. 5a-b)^29^. Using light sheet microscopy, we acquired a brain stack of ~200 μm depth at ~1 volume per second while simultaneously recording the tail motion in a head-embedded preparation (Extended Data Fig. 5c-d). In order to identify the supraspinal neurons recruited during spontaneous locomotion in the two species, we performed a regression analysis on the fluorescence signal of single neurons using regressors representing swimming and termination of swimming (Extended Data Fig. 5e).

This approach revealed neurons in the hindbrain of DT reliably recruited during the termination of swimming and that may therefore be referred as putative “stop neurons” (Fig. 4 a-b). Such stop neurons responsible for termination of locomotion have been reported in other vertebrates such as tadpole, lamprey and mice^30,21,22^. While the short swim events of ZF make it difficult to survey such functional cell types, DT’s long swim events offered a unique opportunity to resolve neurons active at the termination of swimming, and pinpoint motor areas recruited for movement termination. This demonstrates the benefit of employing related animal species in functional studies in addition to their obvious use in understanding evolution of neuronal circuits.

**Fig. 4:**
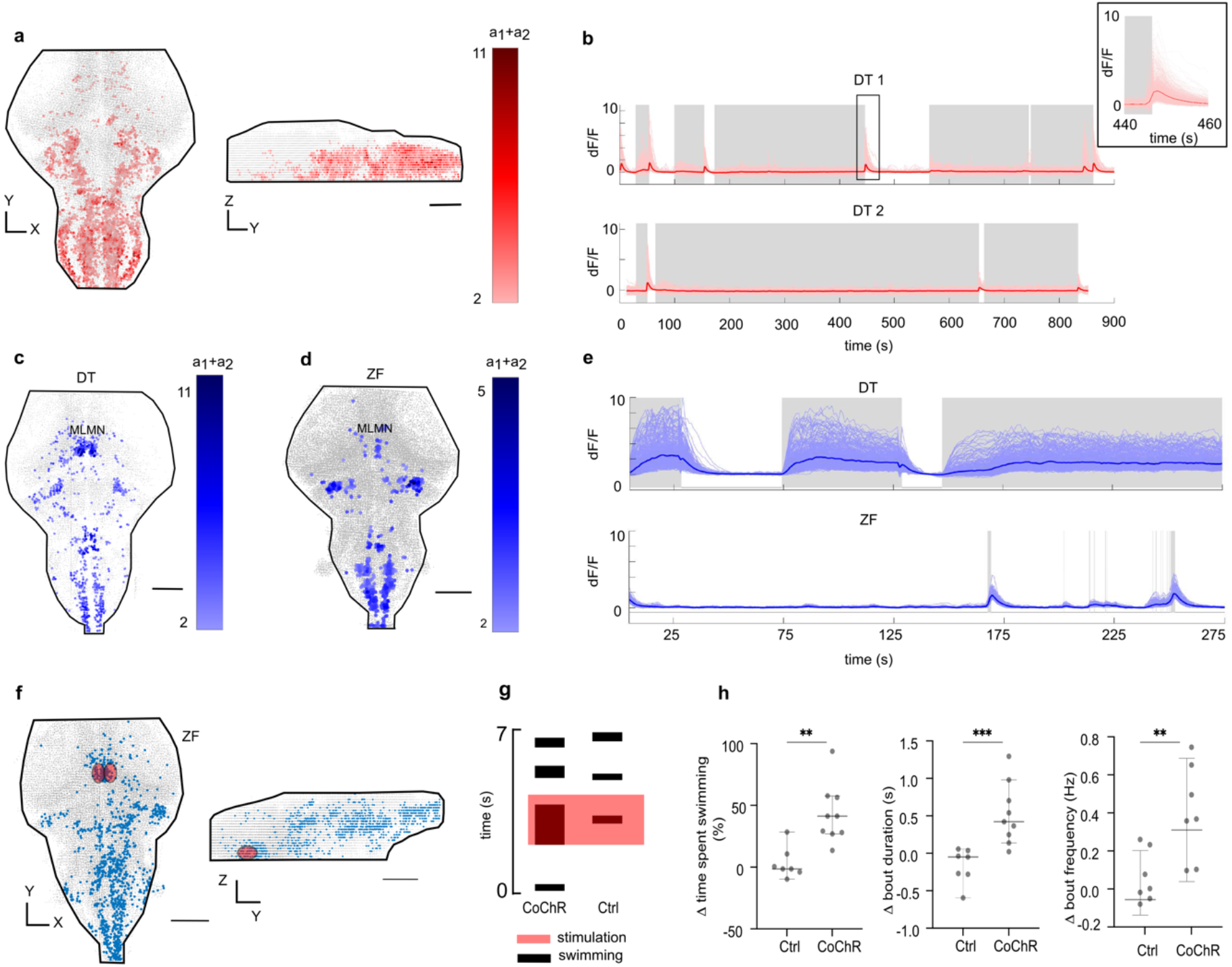
Neurons correlated with termination of swimming (in DT) and swimming (in DT and ZF); and holographic stimulation of the identified Mesencephalic Locomotion Maintenance Neurons (MLMN). (a) Representative images of stop cells (in red) identified in the DT hindbrain (N=3). (b) represents the activity of the identified stop neurons (in red) with respect to swimming activity (in grey). The inset shows a magnified region around a swim termination event. (c) and (d) show a representative figure of a maximum projection of neuronal correlates of swimming (in blue) identified in the DT and ZF brain, respectively. The nuclei correlated with swimming appear to be conserved between DT (N=4) and ZF (N=4). The MLMN population is labelled. (e) Neuronal activity of all the swimming correlated neurons in DT compared to the corresponding neurons’ activity in a ZF for a duration of 300 seconds. The activity in the conserved nuclei in DT is sustained for long durations unlike ZF, correlating with their long swim events. The grey shaded regions represent active swimming. (f) The region of interest (ROI) for holographic optogenetic stimulation is illustrated. MLMN population described in (c) and (d) was first anatomically located under a 2-photon microscope using expression of CoChR-GFP (in test) or GCaMP6 (in control) as the guidance cue. A holographic stimulation protocol was then employed in this ROI (see Extended Data Fig. 7). (g) illustrates the result of the stimulation in a CoChR and Ctrl fish. Swimming activity in CoChR fish is increased during the holographic stimulation. (h) Change in total swimming time, bout duration and bout frequency during optogenetic stimulation. The time spent swimming is prolonged in test fish by the optogenetic stimulation; this increase is caused by both, an increased recruitment of bouts and an increase in the duration of the bouts. N = 7 Ctrl ZF and 9 CoChR ZF. ** p<0.01, *** p=0.001, Mann-Whitney test. All error bars show 95 % confidence interval. Scale bars are 100 μm.

With respect to initiation and maintenance of swimming activity, we identified putative locomotor regions in the brain of larval DT and ZF that were highly correlated with the swim events (Fig. 4c-d, Extended Data Fig. 5). The neuronal activity in these nuclei correlated with the duration of swim events in DT and ZF (Fig. 4e), suggesting a role in the start and / or maintenance of swimming. Among these regions, we identified a strongly correlated midbrain nucleus referred to here as the Mesencephalic Locomotion Maintenance Neurons (MLMN). We decided to focus on this region as the most interesting candidate to sustain the long swim events in DT as previous work in ZF has revealed an anatomically corresponding region suggested to comprise of nMLF as well as other glutamatergic neurons which are implicated in swimming activity^31,32,33^. The nMLF specifically is known to have projections to the caudal hindbrain and spinal cord and plays an important role in locomotor control^31,34^. To dissect the role of MLMN in maintenance of the long swim events, we carried out optogenetic stimulation in ZF *Tg(elavl3:CoChR-eGFP)* which express the opsin CoChR under a pan-neuronal promoter. We targeted MLMN using 2-photon holographic stimulation with temporal focusing (Fig. 4f)^35^ and observed a reliable increase in swimming following the stimulation (Fig. 4g-h, Extended Data Movies 4 and 5). MLMN stimulation led to an increase in the mean duration of bouts and an increase in the frequency of bouts (Fig. 4h), indicating an important role for MLMN in the maintenance of long swim events. The recruitment of swimming on mere stimulation of MLMN alone also suggests that this neuronal population also comprises of some initiation neurons. Interestingly, the discrete swimming patterns of ZF was maintained despite sustained stimulation of the MLMN, suggesting that this property is embedded downstream, possibly in the spinal cord of ZF as previously observed for spinalized ZF preparations deprived of supraspinal inputs^36^. It remains to be further investigated how the long-lasting neuronal activity is produced in DT. The answer may lie in the intrinsic membrane or network properties of the identified neurons and warrants further investigation^37,38^.

In conclusion, using two closely related fish species, we show that two anatomically similar brains with conserved features are able to produce different behavioral outputs based on functional differences in a subset of neurons called MLMN. We also suggest selective pressures which could have led to the divergence of the swimming pattern. This lays the foundation for future work to directly compare neuronal circuits and behaviors in vertebrate species, applying an approach that has been very successfully used in various invertebrate studies^28,39^. This is particularly interesting to perform in danionin fish as many related species are known and can be raised in a laboratory setting. With the ability to assign behavioral modules to their corresponding genetic and neuronal circuit components in ZF (and other danionins), our work provides a powerful approach in comparative neuroethology to investigate evolution of behaviors and neuronal circuits in vertebrates.

## Methods

### Fish breeding and husbandry

6 days post-fertilization (dpf) zebrafish (ZF) and *Danionella translucida* (DT) larvae were used for behavioral experiments. For imaging experiments, 5 dpf ZF and DT were used. DT adults were grown at a water temperature of 25 to 28° C, pH of 6.3 to 8.3 and a conductivity of 250 to 450 uS. Adult DT are fed with Gemma Micro 150 (Skretting, USA) (twice a day) and live *Artemia* (once a day). DT are known to spawn in crevices^3^. Hence, 2 to 4 silicone tubes (~ 5 cm long) were added in the adult tanks to aid spawning.

The larvae were grown at a density of <50 larvae per 90 cm Petri plate in E3 egg medium (without methylene blue). For behavioral experiments, at 5 dpf, both larval ZF and DT were transferred to a 250 ml beaker with 100 ml E3 egg medium (without methylene blue) and fed with rotifers. They were maintained at 28°C in an incubator until the experiment at 6 dpf. The DT larvae were more delicate and required careful handling. Resultantly, the number of DT required to perform each experiment was much larger compared to ZF.

All animal procedures (ZF and DT) were performed in accordance with the animal welfare guidelines of France and the European Union. Animal experimentations were approved by the committee on ethics of animal experimentation at Institut Curie and Institut de la Vision, Paris.

### Free-swimming behavioral acquisition, fish tracking and tail segmentation

A high-speed camera (MC1362, Mikrotron-GmbH, Germany) and a Schneider apo-Xenoplan 2.0/35 objective (Jos. Schneider Optische Werke GmbH, Germany) were used to carry out the free-swimming acquisitions. The resolution of the images were 800 x 800 pixels with 17 pixels /mm and the acquisition was carried out at 700 Hz. The fish were illuminated with an infrared LED array placed below the swimming arena. An 850 nm infrared bandpass filter (BP850-35.5, Midwest Optical Systems, Inc.) was used on the objective to block all the visible light.

The behavioral arena was illuminated with visible light at 220 lux which was similar to the light intensity in the home incubator. The fish were transferred to 35 mm Petri plates and acclimatized for > 2 hours before the behavioral acquisitions. 23 DT and 37 ZF were tested with the acquisition lasting for ~ 20 - 30 minutes.

Image acquisition, fish tracking and tail segmentation were performed using a custom-written C# (Microsoft, USA) program. The online tracking of the fish and tail segmentation was carried out as described earlier elsewhere^40^. Briefly, the following method was performed. A background was calculated by taking the mode of a set of frames which are separated in time so that the fish occupies a different position in each frame. This background image was subtracted from each acquired frame. The subtracted image was smoothed using a boxcar filter. A manually selected threshold was used to separate the fish from the background. The fish blob was selected by performing a flood fill starting at the maximum intensity point. The center of mass of this shape was considered the position of the larva. The middle point on a line joining the center of mass of each eye was defined as the larva’s head position. The direction of the tail was identified by finding the maximum pixel value on a 0.7 mm diameter circle around the head position. Then, a center of mass was calculated on an arc centered along this direction. The angles of ~10 tail segments measuring 0.3 mm were calculated. To do this, successive tail segments were identified by analyzing the pixel values along a 120-degree arc from the previous segment. This same algorithm was used for both DT and ZF. The empirically selected threshold to separate the fish from background was different in the two fish.

### Analysis pipeline for free-swim data

Poor tracking was identified using pixel intensities of the tail segments. The lost frames, if any, were identified based on a 32-bit timestamp encoded in the first 4 pixels of all the images. These lost frames are then interpolated and filled with NaN values for the recorded parameters.

Discontinuities in turning when the fish turns from 0 to 360 degree or vice versa were corrected. The raw X and Y coordinates were smoothed using the Savitzky Golay digital filter: in MATLAB (MathWorks, USA), *sgolayfilt* function is used to implement this. A 2^nd^ order polynomial fit was employed on a window size of 21 units (30 ms). Displacement was calculated using these X and Y coordinates of the centroid of the fish.

The measure of tail curvature was used to identify the bouts^40^. The first 8 tail segments were incorporated in the analysis based on the reliability of the tracking as assessed by the raw pixel intensities. The change in the curvature of the tail was emphasized over local fluctuations by taking a cumulative sum of the values along the tail. The differences in tail angles were calculated as we wanted to detect movements. The tail movements were then smoothed using a boxcar filter of size equivalent to ~14.30 ms. The absolute of the segment angles were then convolved into a single curvature measure. A maxima/minima filter of 28.6 ms/ 572 ms was specifically applied to this tail curvature dataset of ZF based on the knowledge of bout and inter-bout durations available to us. An empirically validated cut-off was used on the convolved and smoothed tail curvature measure to identify the starts and ends of swim bouts. It is important to note that in the analysis, only the ‘burst’ phase of ‘burst-and-glide’ swims were identified in ZF.

The 7^th^ tail segment was used for calculating tail beat frequency and maximum tail angle. The trace of the tail segment was smoothed and small gaps (less than 7 ms) in the swim events due to tracking were interpolated. Swim events with larger gaps were eliminated from the analysis. Any identified events shorter than 71.5 ms in length, if present, were discarded as well to avoid artefacts. On the bout-based kinematics, the bout distance, inter-bout duration, mean and maximum speed, maximum tail angle and tail beat frequency were calculated.

### Half beat based kinematics in ZF and DT

A peak-to-peak half cycle was defined as a half tail beat cycle and this was used for the kinematic calculations to be able to compare a similar unit of locomotion between the two fish. In ZF: to identify the half-beats, on every swim bout, the absolute of the tail angles was calculated from the 8^th^ segment of the tail and the peaks of tail angle were identified using the *findpeaks* function in MATLAB. Extended Data Fig. 1a shows this for a swim bout in ZF.

In DT: the trace of the tail segment is smoothed using a Savitzky Golay digital filter function of 3^rd^ order with a window size of 50 ms. Small gaps (less than 7 ms) in the tracking were interpolated. Using *bwlabel* function in MATLAB on a binary matrix of good/ bad tracking, all continuous stretches of good tracking were labelled. From this, only the stretches longer than 35 frames (or 50ms) were selected for further analysis to avoid small tracking artefacts if any. On these identified stretches, half beats were identified as mentioned above for ZF.

On every half beat in both the fishes, the following kinematic parameters were calculated: duration, distance, mean and maximum speed, maximum tail angle and half beat frequency. The 8^th^ tail segment was used for calculating tail beat frequency and max tail angle.

### Head-embedded swimming

A high-speed camera (MC4082, Mikrotron-GmbH, Germany) with a Navitar Zoom 7000 macro lens was used to carry out the head-embedded acquisitions. The resolution of the images were 400 x 400 pixels with 75 pixels /mm and the temporal resolution of the acquisition was 100 or 250 Hz. The fish were illuminated with an infrared LED array placed below the swimming arena. An 850 nm infrared bandpass filter (BP850-35.5, Midwest Optical Systems, Inc.) was used on the objective to block all the visible light.

6 dpf DT (n=4) and ZF (n=6) were embedded in 0.5 ml of 1.5% agarose. For ZF, nacre incross fish were used. The agarose covered the head up to the pectoral fins. Each fish was acclimatized for at least 90 minutes before acquisition. Recordings lasted for 10-20 minutes per fish. The head-embedded videos were primarily used to determine the amount of time spent in swimming. The tail tracking was performed manually to identify the swimming and resting time periods. The duration of swimming was normalized to the total length of the acquisition and reported as a percentage of the total duration of acquisition.

### Tap-induced escape behavior

An Arduino controlled solenoid was added to the free-swimming behavioral set-up. The Arduino was triggered from the image acquisition program written in C# (Microsoft, USA). When triggered, the solenoid would hit the surface of the arena from the bottom and cause the fish to escape in response to this stimulus. The trigger was only initiated if the fish was not at the edges of the Petri dish and if there was an inter-stimulus interval of at least 50 seconds between two consecutive trials. The delay between the trigger onset and the delivery of the solenoid on the arena was estimated and incorporated in the analysis to calculate an accurate reaction time. 19 DT (n=141 trials) and 15 ZF (n=159 trials) were tested in the assay. Acquisition at 700 Hz was used for the analysis. However, the illustrated images were captured at 100 Hz.

To analyze the escape kinematics, the peak escape velocities were identified in a window of approx. 450 ms after the stimulus delivery. A peak speed was considered as at least 2 times the peak speed during free-swimming (9.25mm/s and 42.5 mm/s for DT and ZF, respectively). In case of multiple peak escape velocities in the window, only the first one was considered. Now a 140 ms region of interest was selected around the peak speed to include 40 ms before the peak and 100 ms after the peak as shown in Extended Data Fig. 2 for a ZF. The region of interest was empirically decided after exploring many trials across both the fish species. Mean speed, total distance covered and the delay to reach the peak speed after the stimulus delivery – these parameters were computed for all the trials in each fish.

The major differences in the processing pipeline from the free-swimming analysis pipeline were as follows. The X/Y displacement vectors were further filtered using a zero-phase digital filtering (*filtfilt* function in MATLAB) with a filter size of 11 ms to identify the peak escape velocities. Kinematics were neither calculated on half beats nor bouts, but on the custom defined 140 ms window for a better comparison of the escape events in the two species of fish.

### Mean squared displacement (MSD) and reorientation analysis

Information on X/Y- coordinates was used to compute the Mean squared displacement (MSD) and decorrelation in heading orientation (R) over time. A Savitsky-Golay filter was applied on the X and Y traces to fit a 2^nd^ order polynomial on a 200 ms window. The filtered trajectories were then downsampled to 70 Hz. For each fish, discrete continuous trajectories were identified in a circular region of interest of radius 18mm to mitigate border-induced bias. These trajectories were used for the computation.

The time-evolution of MSD and R were calculated at every 100 ms time-step and averaged over all trajectories for each animal. To compute R, we extracted at each time t a unit vector **u**(t) aligned along the fish displacement [dx,dy] calculated over a 1s time window. Notice that this vector was only calculated if the fish had moved by at least 0.5 mm in this time period. The heading decorrelation over a period Δt was then computed as R(Δt)=<**u**(t).**u**(t+ Δt)>t. This function R quantifies the heading persistence over a given period: R=1 corresponds to a perfect maintenance of the heading orientation, whereas R=0 corresponds to a complete randomization of the orientations. The MSD and R values were plotted over time for DT (n=23 fish) and ZF (n=37 fish).

### Quantification of depth preference

Three vertical glass cylinders with 36 cm water height were used in this experiment. 6 dpf ZF larvae (n=30 per cylinder) were added to three cylinders. 6 dpf DT larvae (n=30 per cylinder) were added to another three cylinders.

The cylinders were considered as consisting of three sections and marked accordingly: the bottom 12 cm, middle 12 cm and the top 12 cm. The number of fish in each section of the column was manually counted once every hour for 10 hours. Only the fish that were swimming normally were considered for the enumeration. This was used to calculate the average normalized fish density in every section of the water column.

### Quantification of body length and swim bladder inflation

Body length was measured in 5 dpf larvae of the two species (n = 10 for DT and n = 9 for ZF). Pictures of the larvae were captured using an AxioCam MR3 camera. The magnification of the optics was noted and the physical dimension of the camera pixel was used to calculate the pixel size in μm as follows: pixel size = (260/magnification) x binning factor.

To quantify swim bladder inflation, from a population of growing larvae (3 dpf to 15 dpf), five or more larvae were sampled for each age and the proportion of larvae with inflated swim bladder was quantified. The sampling was performed from a growing population of approximately N = 30 DT and 30 ZF.A moving averaging was performed using a window size of two units to smoothen the curve and the swim bladder inflation results were reported from 3.5 to 14.5 days.

### Whole-mount *in-situ* hybridization (ISH)

To generate anti-sense probes, DNA fragments were obtained by PCR using Phusion™ High-Fidelity DNA Polymerase (Thermo Scientific™) and the following primers (5’->3’)^41^: vglut2a (forward primer: *AGTCGTCTAGCCACAACCTC;* reverse primer: *CACACCATCCCTGACAGAGT*), vglut2b or *slc17a6b* (forward primer: *GCAATCATCGTAGCCAACTTC*; reverse primer: *ACTCCTCTGTTTTCTCCCATC*), glyt2 or *slc6a5* (forward primer: *TGGAAGGATGCTGCTACACA;* reverse primer: *TGACCATAAGCCAGCCAAGA*) and gad67 or *gad1b* (forward primer: CCTTCCTCCTCGGCGATTGA; reverse primer: GGCTGGTCAGAGAGCTCCAA). Total cDNA for ZF and DT were used as a template. PCR fragments were cloned into the pCRII-TOPO vector (Invitrogen) according to manufacturer’s instructions. All plasmids used were sequenced for confirmation.

Digoxigenin RNA-labeled or Fluorescein RNA-labeled probes were transcribed *in vitro* using the RNA Labeling Kit (Roche Diagnostics Corporation) according to manufacturer’s instructions. Dechorionated embryos at the appropriate developmental stages were fixed in fresh 4% paraformaldehyde (PFA) in 1X phosphate buffered saline (pH 7.4) and 0.1% Tween 20 (PBST) for at least 4 hours at room temperature or overnight at 4° C. Following this, the samples were preserved in methanol at −20° C until the *in-situ* experiments described below. Whole-mount digoxigenin (DIG) *in-situ* hybridization was performed according to standard protocols^42^. A protease-K (10 μg/mL) treatment was performed depending on the age and species of the sample (90 minutes and 120 minutes for 5 dpf DT and ZF, respectively). The samples were imaged on a stereoscope with AxioCam MR3 camera.

### Vibratome section

The whole-mount samples were embedded in gelatin/albumin with 4% of Glutaraldehyde and sectioned at 20 μm thickness on a vibratome (Leica, VT1000 S vibrating blade microtome). The sections were mounted in Fluoromount Aqueous Mounting Medium (Sigma) before imaging.

### Whole-mount Fluorescence *in-situ* hybridizations (FISH)

The samples stored in methanol at −20°C were rehydrated by two baths of 50% methanol/PBST followed by two baths of PBST. This was incubated for 10 min in a 3% H2O2, 0.5% KOH solution, then rinsed in 50% methanol/ 50% water and again dehydrated for 2 hours in 100% methanol at −20°C. Samples were rehydrated again by a series of methanol baths from 100% to 25% in PBST, and washed two times in PBST. This was followed by an age and species dependent treatment of proteinase-K (10 μg/mL) at room temperature. At 5 dpf, DT and ZF underwent treatment of proteinase-K for 90 and 120 minutes, respectively.

Following this, the samples were again fixed in 4% PFA/PBST. After 2 hours of pre-hybridization in HY4 buffer at 68° C, hybridization with fluorescein-labelled probes (40ng probes in 200 μl HY4 buffer) was performed overnight at 68°C with gentle shaking. Embryos were rinsed and blocked in TNB solution (2% blocking solution (Roche) in TNT) for 2 hours at room temperature. This was then incubated overnight with Fab fragments of anti-Fluo-POD (Roche) diluted 1:50 in TNB. For signal revelation, embryos were washed with 100 μl Tyramide Signal Amplification (TSA, PerkinElmer) solution and incubated in the dark with Fluorescein (FITC) Fluorophore Tyramide diluted 1:50 in TSA. The signal was then followed for 30 minutes for *glyt* and 1 hour for *vgult2b* until a strong signal was observed. After which, the reactions were stopped by 5 washes with TNT, and incubated for 20 minutes with 1% H2O2 in TNT. All the steps after Fluorescein (FITC) incubation were processed in the dark.

### Immuno-histochemistry

Briefly, the whole mount embryos were washed twice in TNT solution. Subsequently, they were blocked for 1 hour at room temperature in 10% Normal Goat Serum (Invitrogen) and 1% DMSO in TNT solution. Rabbit GAD65/67 primary antibody (AbCam) diluted 1/5000 in 0.1% blocking solution was incubated overnight at 4°C. The Alexa Fluor 635 secondary antibody goat anti-rabbit IgG (1/500) (Life Technologies) was added in 0.1% blocking solution and incubated overnight at 4°C. After 5 washes in PBST buffer, microscopic analysis was performed.

### Confocal imaging of the whole brain FISH/IHC samples

To image the whole brain *in-situ* hybridization and immunohistochemistry samples, we used Zeiss LSM 780, LSM 800 and LSM 880 confocal microscopes with a 10x or 40x objective using appropriate lasers and detection schemes suitable to the labelled sample. Whole brain images were acquired in tiles and stitched together using the stitching algorithm available in Zeiss ZEN blue and ZEN black. The images are shown as maximum intensity projections created on imageJ. In GAD65/67 IHC, non-specific blobs of signal likely originating from residual dye left on the skin after the washing step was removed using image processing in the representative image.

### Retrograde labelling of reticulospinal (RS) neurons

A solution containing 10% w/v Texas Red dextran (TRD, 3,000 MW, Invitrogen) in water was pressure injected in the spinal cord (between body segment 7 to 14) of 4dpf ZF and DT. In DT, this method resulted in less efficient labeling of the RS neurons. The best results were obtained by cutting the tail beyond segment 14th with fine scissors and pressure injecting the TRD in the exposed spinal cord. After the labeling, the fish were allowed to recover in E3 egg medium for 24 hours at 28° C.

At 5dpf, the surviving injected larvae were anaesthetized with 0.02% Tricaine (MS-222, Sigma), mounted in 1.5% low melting point agarose and imaged under a VIVO 2-photon microscope (3I, Intelligent Imaging Innovations Ltd). Labelling was often sparse and varied among the injected fish which survived to 5dpf (n=4 fish per species). Maximum intensity projection images of the reticulospinal neurons in ZF and DT is shown from the animals where almost all the RS neurons were labelled. RS neurons in the DT brain were annotated based on their anatomical similarity to the ones in ZF^23^.

### Generation of pan-neuronal calcium sensor *Tg(elavl3-H2B:GCaMP6s)* line

To generate *Tg(elavl3:H2B-GCaMP6s)* DT fish, 6 ng/μl of the plasmid and 25 ng/μl of Tol2 was used. Injections were performed in embryos which were less than or equal to 4-cell stage. The injection was performed free-hand as DT lay eggs in clutches. Tol2-elavl3-H2B-GCaMP6s plasmid was a gift from Misha Ahrens (Addgene plasmid # 59530; http://n2t.net/addgene:59530; RRID: Addgene_59530)^44^.

### Light sheet microcopy

Transgenic DT and ZF expressing H2B-GCaMP6s under the *elavl3* promoter were utilized. The GCaMP is nuclear tagged, so its expression is limited to the nucleus which makes it easier for segmentation of the neurons. The fish were embedded in a capillary with 2.5% agar. The tail was freed and recorded simultaneously to extract a readout of the spontaneous swimming behavior. Before each recording, the embedded fish were acclimatized to the recording chamber with the blue laser switched on for at least 10 minutes. The scanning objective was lateral and the beam entered from the left side of the fish and the detection objective was placed upright on the top. Both the objectives were moved with a piezo so that the light sheets were always in the focal plane of the detection objective. Average laser power was at 0.05 mW. Approximately 280 μm of the brain volume was imaged in each fish. Brain imaging was carried out at approximately 1 Hz (1 whole brain volume / s) and the tail movement was acquired at ~ 40-80 Hz. Each acquisition lasted for ~ 20 minutes.

### Image processing and analysis pipeline for whole-brain light-sheet data

For ZF and DT, image processing was performed offline using MATLAB. Based on visual inspection, if needed, image drift was corrected by calculating the cross-correlation on a manually selected region of interest (ROI). The dx and dy values employed to correct the drift in this ROI were extrapolated to the whole stack. Brain contour was manually outlined on mean greystack images for each layer. Background value for each layer was estimated from the average intensity of pixels outside the brain contour. The segmentation procedure consisted of a regression with a Gaussian regressor convolved with the same Gaussian regressor. The result was regressed another time with the same regressor. Baseline and fluorescence were calculated for each neuron identified by the segmentation. The fluorescence *F(t)* signal was extracted by evaluating the mean intensity across the pixels within each neuron. The tail tracking was performed manually on the tail acquisitions to identify active and inactive time periods. For ZF, the baseline was calculated by the running average of the 10th percentile of the raw data in sliding windows of 50 seconds. For DT, the baseline for identifying neurons correlated with swim and stop events was calculated as the 10th percentile of the raw data within each inactive period defined as the time period between 5 seconds after the end of a swim event and 3 seconds before the beginning of the next swim event (from tail acquisition data). The baseline values for the active periods were interpolated using the values in the inactive periods. For both ZF and DT, the relative variation of fluorescence intensity *dF/F* was calculated as *dF/F* = *(F(t)* – baseline) / (baseline-background).

For both ZF and DT, neurons from the more rostral part of the brain were removed (y coordinates between y_max_ – 10 um and y_max_) because of dF/F artefact due to image border. A multi-linear regression was performed using the classical normal equations. In DT, this was performed on dF/F for the whole duration of the experiment and in ZF, on dF/F for a manually selected time period with many well isolated swim bouts. The analysis determines the best-fit coefficient β to explain the neuronal data (y) by the linear combination *y = Σβ_j_ * x_j_* + *β_0_*, where *x_j_* is the regressor. For ZF, a constant regressor and a swim maintenance regressor (based on the tail acquisition data) were used. For DT, four regressors were used: constant, swim maintenance, swim onset and swim offset. The onset and offset regressors were obtained from the initiation and termination of swim events (based on the tail acquisition data) with a time window of −3 seconds to +1 second around the initiation/ termination event. Swim maintenance, onset and offset regressors were convolved with a single exponential of 3.5 seconds decay time which approximates the H2B-GCaMP6s response kernel in ZF^45^. T-scores were computed for every neuron/regressor combination. We could reliably find neurons highly correlated with swim maintenance and termination events as shown in the results.

### Brain registration

We used the Computational Morphometry ToolKit CMTK ((http://www.nitrc.org/projects/cmtk/) to compute and average the morphing transformation from high resolution brain stacks (184 layers and 1μm z-resolution; 1-photon imaging) to create a common brain for the *Tg(elavl3:H2B:GCaMP6s)* DT line. All the calcium imaging results were mapped to this reference brain. To compare neuronal populations across different brain samples, we calculated the spatial densities of the considered clusters by using the Kernel Density Estimation (KDE) with a Gaussian kernel with a bandwidth of 12.8 μm. Discrete cluster densities were determined for all points of an inclusive common 3D rectangular grid with an isotropic resolution of 5 μm.

### Optogenetic stimulation

5 to 7dpf ZF were head-embedded in a Petri dish with 2% agarose and the tail was freed to move. After a period of acclimatization, fish were placed under a custom made 2-photon (2P) microscope capable of 2P scanning imaging and 2P holographic patterned illumination^46^. The holographic optical path is analogous to the one described in a previously published work^47^. Briefly, the use of fixed phase mask, a diffraction grating and liquid crystal spatial light modulator allows the generation of multiple illumination spots distributed in 3 dimensions^47^. Additionally, an inverted compact microscope and an infrared LED (780 nm) were placed below the Petri dish to record the tail movements. 2P scanning imaging of *Tg(elavl3:CoChR-eGFP)* and *Tg(elavl3:H2B-GCaMP6s/6f)* (control) was first performed to locate the MLMN in the midbrain. To target the MLMN population, we defined a holographic illumination pattern composed of multiple identical holographic spots distributed over different x-y-z locations. Each spot has a lateral diameter of 12 μm and an axial FWHM ≈ 10μm. On the x-y plane, the targeted surface is covered by the generation of 10 holographic spots, then this pattern is reproduced over 3 different planes to adjust the axial extension of the excitation volume (Extended Data Fig.7). The resulting excitation region corresponds approximately to an ellipsoid of 40-50-70 μm (x-y-z axis, respectively), matching the size of the MLMN in each hemisphere (see Fig. 4 f). Neurons in these regions were photo-stimulated by 2P excitation with the following protocol: 10 ms pulses at 10Hz were delivered for 2 s and repeated 3 times with 30 s intervals between repetitions. The effective excitation light intensity varied from 25 to 40 μW/μm^2^ and was delivered through a 40x objective (N40X-NIR,0.8 NA,Nikon) by an amplified fiber laser at 1040 nm (Satsuma,Amplitude System), suitable to efficiently excite CoChR opsin^48^. Simultaneous recording of the tail movement was performed on a CMOS camera (MQ013MG-ON Ximea) at a frame rate of 33Hz. For analysis, the tail tracking was performed manually with the respect to the periods of stimulation. We extracted three swimming parameters during both, spontaneous swimming and stimulation protocol: bout duration, bout frequency and proportion of time spent swimming. The increase or decrease in the mean value of these parameters during the stimulation protocol for each animal is represented in Fig. 4h.

## Statistical methods

### Behavior data

All the averaged values per fish were prepared in MATLAB 2017b (Mathworks) and statistical tests between the populations were carried out in Prism 8 (GraphPad). Mann-Whitney test by ranks was performed in all cases where the dataset did not follow a normal distribution.

### Light-Sheet imaging data

To characterize highly responsive neurons for a specific regressor, the regression coefficient and t-score distributions were first fitted with a Gaussian model (μ_dist_, σ_dist_) to estimate a sub-distribution responsible for noise (neurons that do not correlated well with the regressor). These sub-distributions, defined as the maximum distribution ± σ_dist_, were then fitted again with a Gaussian model (μ_nosie_, σ_noise_). The highly responsive neurons were defined as neurons with both, a regression coefficient higher than regression threshold_coefficient_ = μ_noise coefficient_ + 3 σ_noise coefficient_ (or threshold_coefficient_ = μ_noise coefficient_ + 4 σ_noise coefficient_) and a t-score higher than t-score threshold_t-score_ = μ_noise t-score_ + 3 σ_noise t-score_ (or threshold_coefficient_ = μ_noise t-score_ + 4 σ_noise t-score_). To quantify the responsiveness of highly correlated neurons, a score was created for each neuron based on the sum of the regression coefficient normalized by the regression threshold_coefficient_ (a_1_) and the t-score normalized by the t-score threshold_t-score_ (a_2_). The higher the score, the more responsive is the neuron.

## Acknowledgements

We would like to thank Mykola Kadobianskyi for providing us with an early access to the *Danionella translucida* genome. Thanks to Adrien Jouary and Michael Orger for sharing their expertise in animal tracking and Roshan Jain for helpful discussion. G.R. was supported by a European Union’s Horizon 2020 research and innovation programme under the Marie Skłodowska-Curie grant agreement No 666003, a Fondation pour le Recherche Medicale (FRM) 4th year doctoral fellowship and a Sorbonne University Postdoctoral Fellowship. Work in the laboratory of F.D.B. was supported by ANR-18-CE16 “iReelAx”, UNADEV in partnership with ITMO NNP/AVIESAN (national alliance for life sciences and health), the Fondation Simone and Cino del Duca the Programme Investissements d’Avenir IHU FOReSIGHT (ANR-18-IAHU-01). B.J. acknowledges support by the German Research Foundation (DFG, project EXC-2049-390688087 and project 432195732), the Einstein Foundation (EPP-2017-413), the European Research Council (ERC-2016-StG-714560) and the Alfried Krupp Foundation. C.G. was supported by an EMBO Short-term Fellowship, an Institut Curie Postdoctoral Fellowship and an FRM Postdoctoral Fellowship. G.D. was supported by H2020 European Research Council (71598).

## Author Contributions

G.R. and F.D.B. conceived the project with inputs from C.W., G.D. and C.G. G.R. designed all the experiments, developed the behavior rig and transgenic fish, and performed all the experiments and analysis unless otherwise specified. J.L. performed the analysis of the whole-brain data with inputs from G.R. under the supervision of G.D. M.C.T. and G.R. performed the backfill experiments under the supervision of C.W. K.D. and G.R. performed the other anatomical experiments. D.T. and G.F. performed the optogenetic experiment and analysis under the supervision of V.E. T.P. and R.C. built the light-sheet imaging rig. J.H., B.J. and R.B. obtained data from the field study. F.D.B. and C.G. supervised G.R. G.D. wrote the MSD/reorientation analysis script. The manuscript was written by G.R. and F.D.B. with inputs from other authors. All authors read and approved the final manuscript.

## Notes

### Competing Interest Statement

The authors have declared no competing interest.

