## Supplemental Material for "Evolutionary divergence of locomotion in two related vertebrate species"

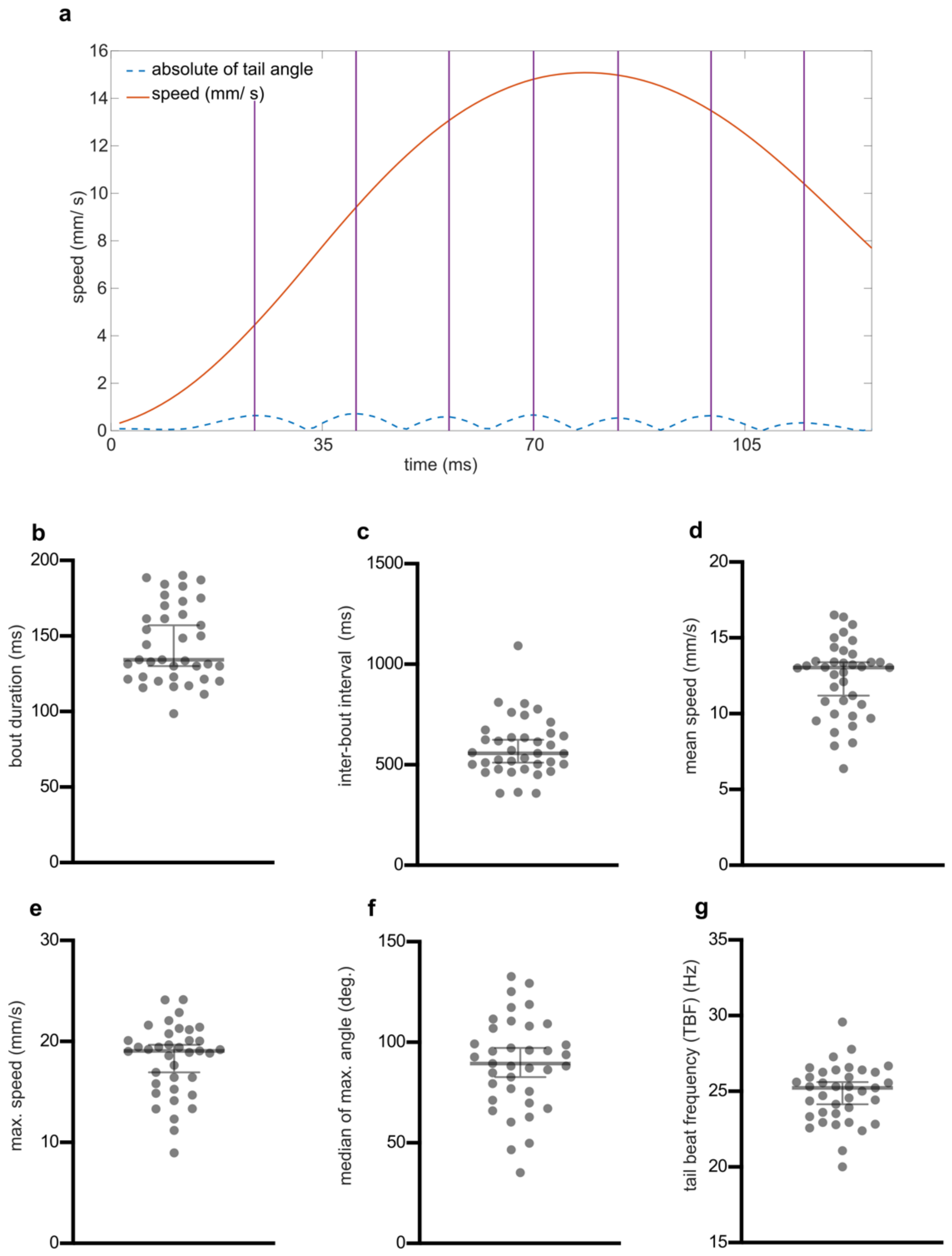


**Figure 1: Half tail-beat detection and bout-wise kinematic analysis.**

(a) Illustration of half tail-beat identification in a swim bout of 6 dpf ZF. The vertical lines show the peak of tail bend which is considered as the start and/or end of a half beat. It is identified based on the peaks of the absolute tail angle (shown in cyan; not to be quantified). These half tail beats are used to calculate the spontaneous swimming kinematics. (b) The median bout duration, (c) inter-bout duration, (d) median of mean speed and (e) maximum speed, (f) median of maximum tail angle and (g) tail-beat frequency (TBF) of ZF after analyzing the same dataset (as used in the half beat based calculations) utilizing a bout-wise analysis method (N=37 ZF each). The values obtained by performing such a bout-wise analysis were found to be close to what has been reported for ZF earlier [3].

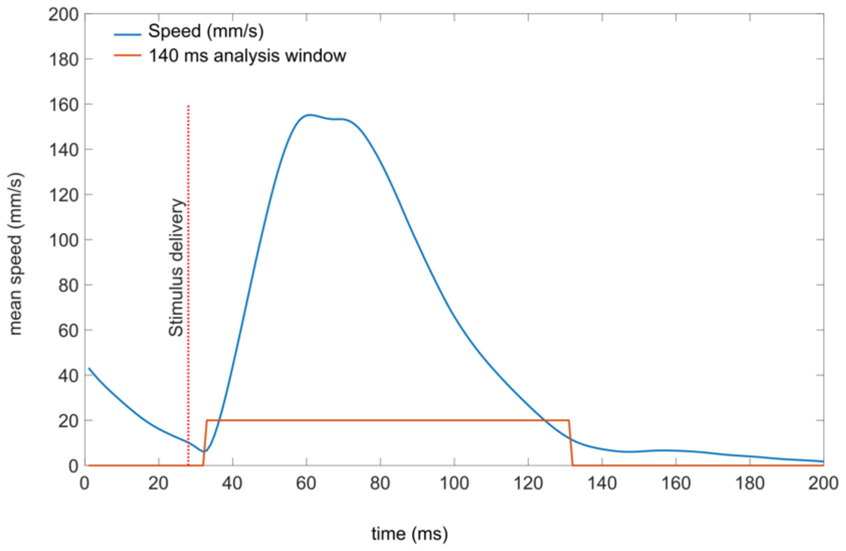


**Figure 2: Analysis of escape kinematics.**

Illustration of the 140 ms window selected for the analysis of tap-induced escape kinematics in ZF. The vertical dotted red line shows the time of stimulus delivery. The region occupied by the horizontal red line illustrates the 140 ms window used for analysis of the escape kinematics.

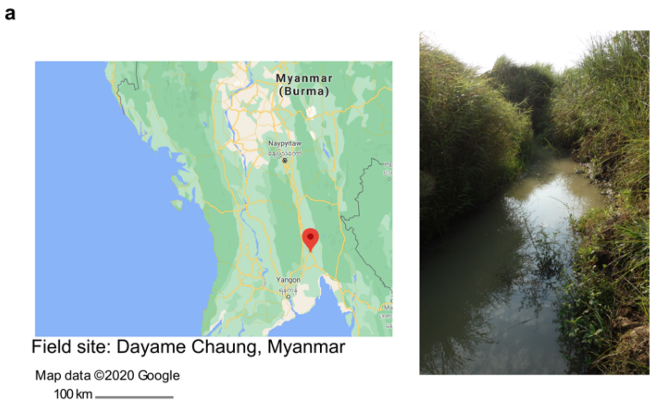


| Depth of sampling point (cm) | $O_2$ Concentration (mg/L) | Temperature (ºC) |
|---|---|---|
| 0 (surface) | 8.75 | 27 |
| 50 | 3.5 | NA |
| 80 | 2.8 | 24 |

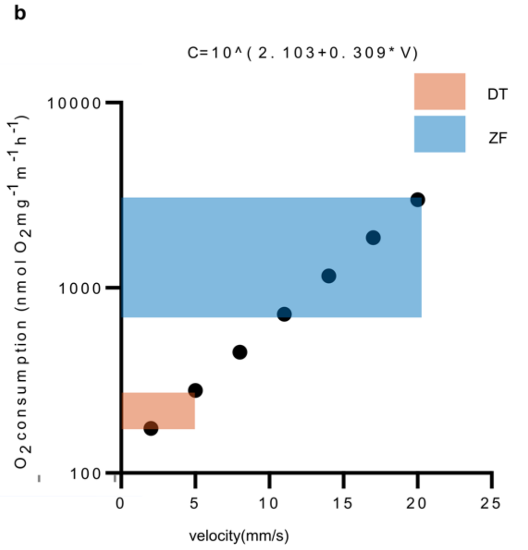


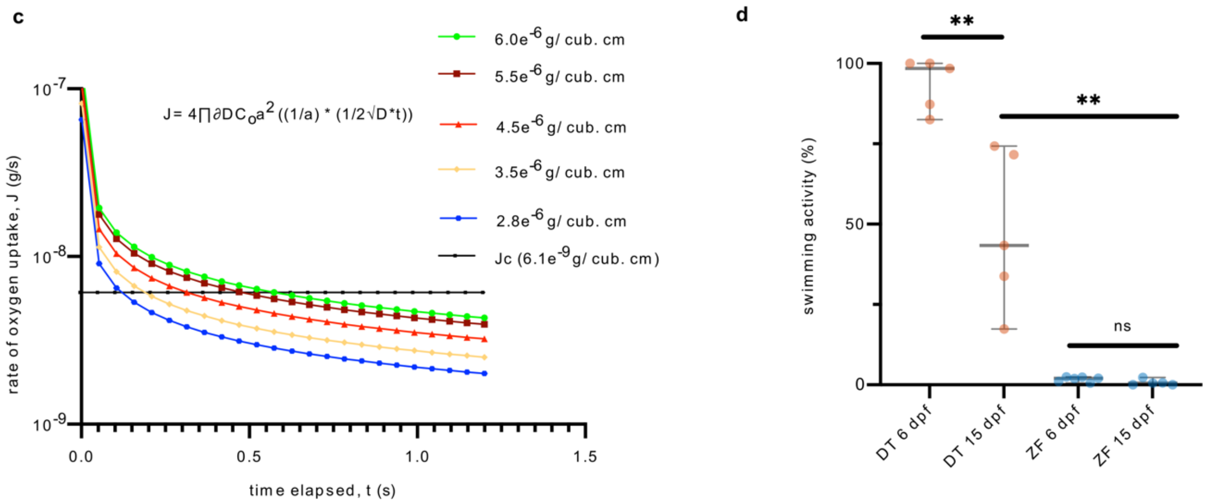


**Figure 3: Influence of oxygen availability and swim bladder on swimming pattern.**

(a) Map (left) shows the site of field collection: Dayame Chaung, Myanmar. Photograph (right) shows the side channel where maximum number of DT were spotted. Table (below) shows the oxygen concentration and temperature found at various depths of the channel at this site. (b) Many studies have looked at the relationship between speed of locomotion and oxygen consumption. Using a relationship derived by Bagatto *et al.* in larval ZF between oxygen consumption and swimming speed, here we try to show how the oxygen demand would vary for a ZF-like larva swimming in the range of speeds reported for DT and ZF[4]. Log C = 2.103 +

0.309 x V, where C is a measure of oxygen consumed per mg of the animal per hour and V is the speed. DT swimming at lower speeds would have a lower oxygen demand compared to ZF.

(c) Analytical model showing lower availability of dissolved oxygen (DO) in the surrounding can lead to a shorter inactivity time between swim events, leading to a more continuous swimming (based on early work of Weihs[5]). The rate of oxygen uptake ($J$) over time ($t$=0, when the animal stops moving) depends on the initial concentration of dissolved oxygen ($C_o$) in the surrounding water (shown in colored curves). $\partial$ is density of water: 1 gm/cm$^3$, $D$ is diffusion constant for dissolved oxygen at 22°C: 2.22E-5 and a is the radius of an equivalent sphere, the surface area of which is equal to the surface area of the larval body. Horizontal line ($J_c$) indicates the approximation of the critical oxygen uptake rate, beyond which the larvae would have to start to swim again in order to replenish oxygen in its surrounding (calculated based on the inter-bout duration of ZF). From the plot: as the $C_o$ decreases, the $J$ drops more rapidly; and this leads to a lower delay to reach the $J_c$, giving rise to more continuous DT-like swimming with shorter pauses. The range of DO values used is based on the values recorded for DT and ZF in nature[6]. (d) Percentage of time spent swimming in 15 dpf DT with swim bladder (n=5), 6 dpf DT without swim bladder (n=5), 6 dpf ZF with swim bladder (n=6) and 15 dpf ZF with swim bladder (** $p<0.01$, Mann-Whitney test). Swimming activity in 15 dpf DT larva is decreased when compared to 6 dpf DT larva. However, the swimming activity in 15 dpf DT is still significantly higher than ZF at both 6 and 15 dpf with swim bladder.

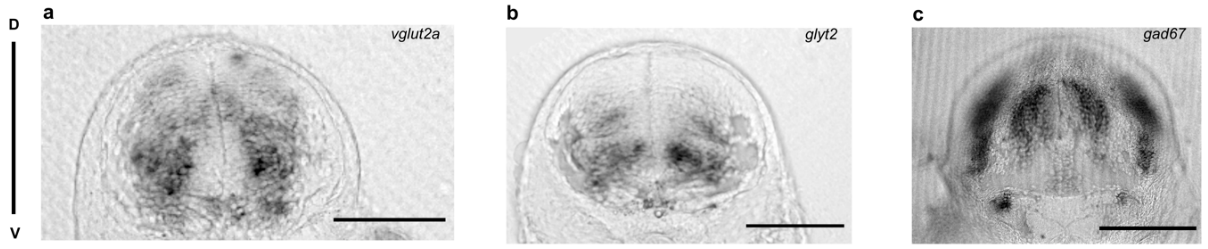


**Figure 4: DT hindbrain cross-section showing glutamatergic, glycinergic and GABAergic neurons.**

Representation of cross-sectional view of DT hindbrain showing distribution of (a) *vglut2a* (or *slc17a6b*), (b) *glyt2* (or *slc6a5*) and (c) *gad67* (or *gad1b*) markers following digoxigenin (DIG) *in-situ* hybridization. Scale bars are 50 µm.

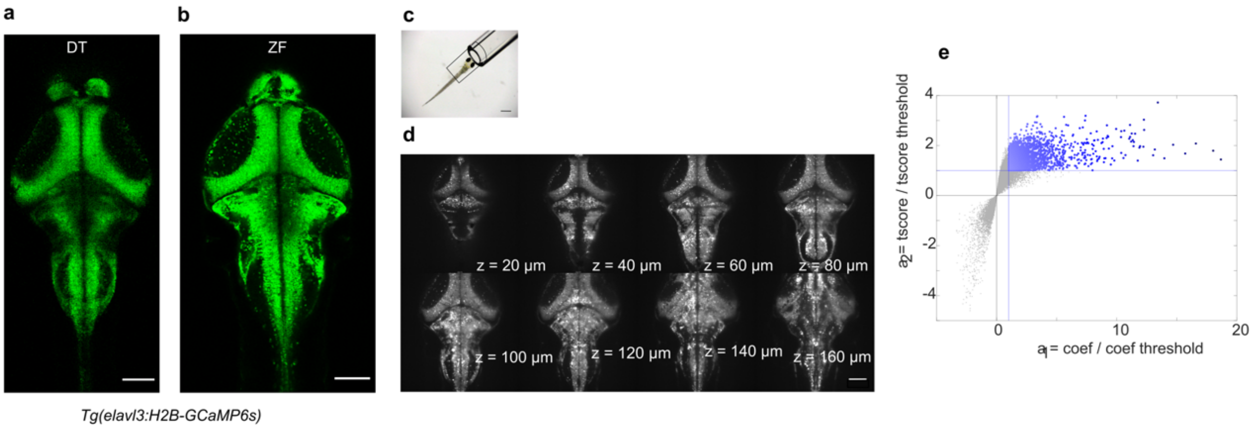


**Figure 5: Light-sheet imaging with 5 dpf *Tg(elavl3:H2B-GCaMP6s)* DT and ZF.**

(a) A maximum intensity projection of 5dpf *Tg(elavl3:H2B-GCaMP6s)* DT. (b) A maximum intensity projection of 5 dpf *Tg(elavl3:H2B-GCaMP6s)* ZF. The expression pattern of the GCaMP appears very conserved between the two larvae. Scale bars are 100 µm. (c) Holding capillary to immobilize the fish during imaging. 2.5% agar is used to hold the fish in the capillary. 1/3$^{rd}$ of the body (tail-region) is freed from the agar. The tail activity is recorded during the brain imaging and used as readout for the swimming activity.

(d) A 5dpf DT brain at various depths represented by Z (0 µm being most dorsal). Approximately 200 µm of brain (in 8 µm steps) in DT and ZF was acquired at 1 Hz during simultaneous recording of tail activity to identify and compare the activity of neuronal correlates of locomotion in the two species. Scale bar is 100 µm. (e) An illustration of the selection of swim maintenance cells based on their regression coefficient ($a_1$) and t-score ($a_2$). The neurons are encoded in a color gradient based on their ($a_1$+ $a_2$) score. The color gradient increases as the score increases.

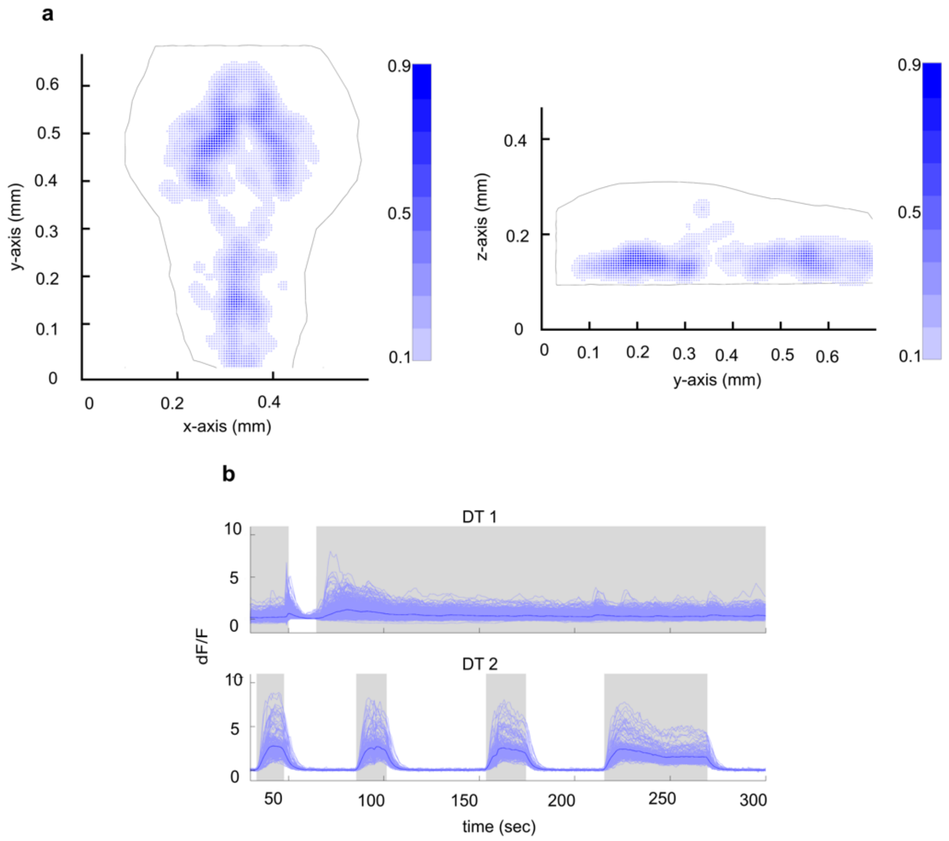


**Figure 6: Neuronal correlates of swimming in DT.**

(a) A KDE (kernel density estimator) plot showing the probability of a neuron in DT brain to be a swim maintenance neuron. The KDE is estimated based on regression data from N= 4 DT brains. (b) Neuronal activity of all the swimming correlated neurons in two DT fish for a duration of 300 seconds. The activity in the nuclei are sustained for long durations correlating with the long swim events. The grey shaded regions represent active swimming.

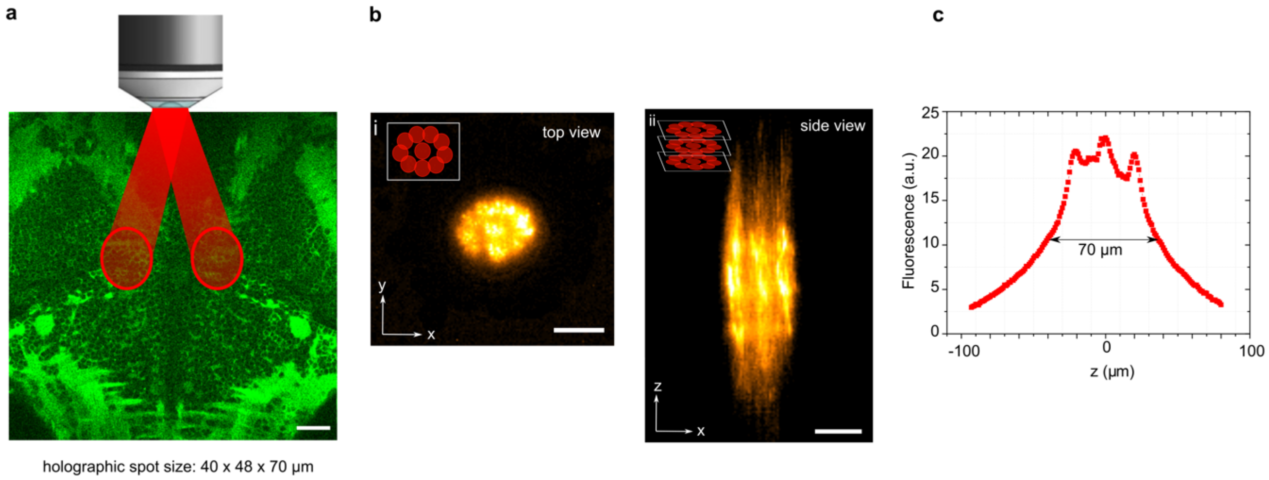


**Figure 7: Two-photon holographic illumination of MLMN.**

(a) An illustration of the two-photon holographic illumination. (b) Top-view (left) and lateral view (right) of the excitation volume of the holographic spot used to target the MLMN population in each hemisphere. Laterally, the targeted surface is covered by the generation of multiple temporally focused holographic spots; each spot with a circular shape of 12 µm diameter (inset i). This x-y pattern is then reproduced over 3 different planes generating an axially extended excitation volume (inset ii). The corresponding integrated axial profile of the spot is reported in (c), showing a FWHM of ≈ 70 µm, matching the z-extension of the targeted population. Scale bars are 30 µm.

**Movie 1:** Movie of a 6 dpf *Danionella translucida* (DT) exploring its environment.

**Movie 2:** Movie of a 6 dpf zebrafish (ZF) exploring its environment.

**Movie 3:** Assay used to study the tap-induced escape response.

**Movie 4:** Tail movement during optogenetic stimulation of a *Tg(HuC:CoChR-eGFP)* ZF.

**Movie 5:** Tail movement during optogenetic stimulation of a *Tg(HuC:H2B-GCaMP6s)* ZF (control).
